# Analysis of 3’-seq data from multiple *E. coli* studies identifies diverging results sets and raw data characteristics despite similar collection conditions

**DOI:** 10.1101/2025.06.12.658996

**Authors:** Quinlan Furumo, Michelle M. Meyer

## Abstract

3-prime end sequencing (3’-seq) is a high-throughput sequencing technique that is used to specifically quantify the changes in 3’-end formation of transcripts in bacterial cells, which is increasingly being utilized to address fundamental questions regarding transcription termination and pausing across a range of different bacterial species. However, the growing number of 3’-seq studies is accompanied by an increase in study-specific 3’-seq data analysis approaches. Thus, differences in a number of factors including: experimental design, data collection approaches, analysis methodologies, and interpretation decisions, make it challenging to confidently compare results derived from different studies, even those that were performed on the same organism. To assess the potential severity of these discrepancies, we used PIPETS, a statistically robust and genome-annotation agnostic 3’-seq analysis package, to study *Escherichia coli* 3’-seq data sets from three different groups collected under similar conditions. By using a consistent analysis and results interpretation approach, we identified large disparities in the characteristics of the raw 3’-seq data between each of the studies, despite all three studies using the same strain and very similar reported experimental conditions. Additionally, we found strand-specific inconsistencies, with some data sets having reference strand 3’-seq read coverage distributions that differed greatly from the complement strand within the same replicate. Finally, when the 3’-seq distribution profiles of the three *E. coli* studies are compared to studies from four additional bacteria, we identified 3’-seq results clustering patterns that are not explained by phylogenetic similarity between organisms. With the large differences seen between data sets from the same organism as well as the inconsistencies seen between replicates from the same data sets, we urge the field to reconsider the assumptions around 3’-seq data homogeneity and move towards consistent analysis approaches, and cautious interpretation of the data.

**Graphical Abstract:** 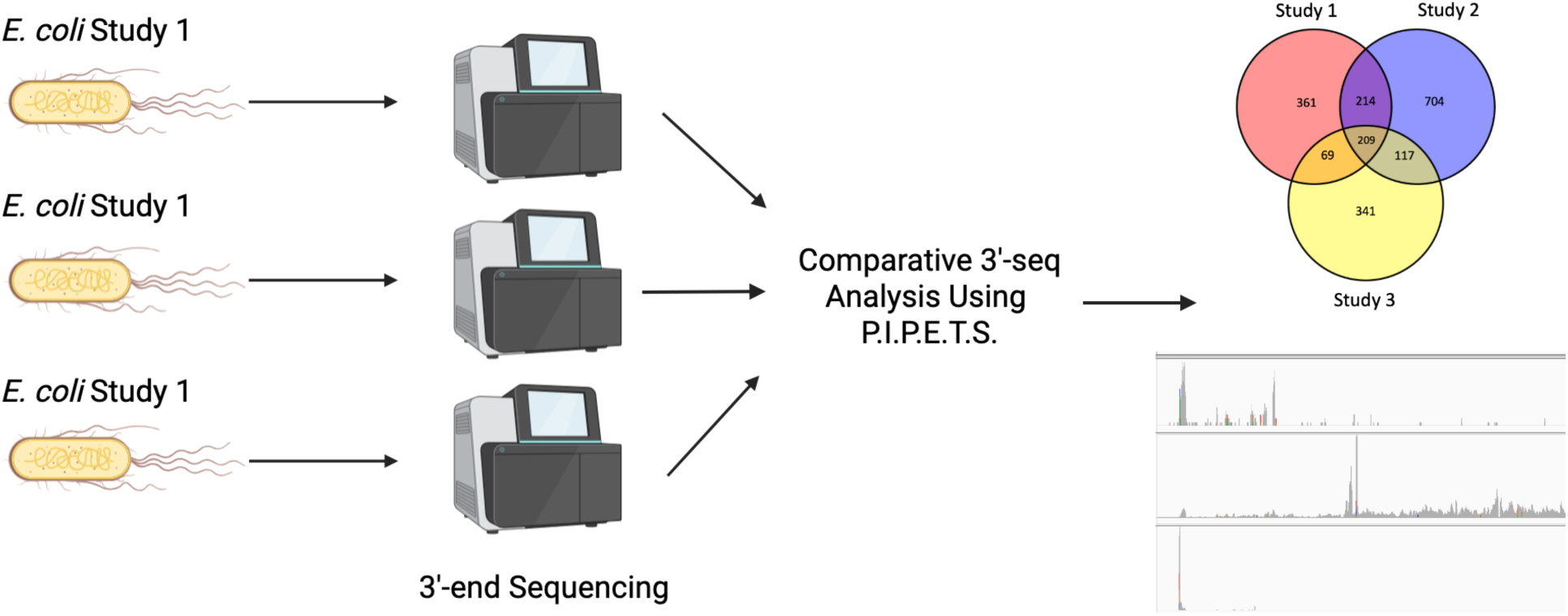

## Introduction

Bacterial transcriptional dynamics is a rapidly growing field of research that provides insight into a myriad of biologically and medically relevant processes of bacteria. This growth is due, in part, to the widespread sequencing and study of transcript 3’-ends using two primary methods: 3’-end sequencing and extrapolated bulk-RNAseq. Term-Seq/3’-end Sequencing (3’-seq) is a high-throughput RNA sequencing method that non-specifically tags the 3’-ends of transcripts for sequencing. In the Gram negative model organism *Escherichia coli*, 3’-seq has been used to discover novel transcriptional regulatory networks that provide a more comprehensive understanding of transcriptional dynamics in both Rho-dependent and Rho-independent contexts^1–3^. Studies using non-traditional model organisms have also effectively used 3’-seq to analyze the abundance and classification of different types of transactional regulation^4–7^, to characterize transcriptomes of non-traditional model organisms alongside additional types of RNA sequencing^8–10^, and to study evolutionary conservation of newly cultured strains and organisms^11,12^. Transcript 3’-ends may also be extrapolated computationally from more traditional RNA-sequencing experiments. For the Gram positive model organism *Bacillus subtilis*, such approaches have been used to study the evolution of antimicrobial resistance^13^, exonuclease activity and prevalence for mRNA decay^14^, and the roles of NusA and NusG in elongation and transcriptional termination^15^. As the number of species and strains being studied using both methods increases, so too does the volume of data available that enables cross-species comparisons that might draw wider scope biological conclusions regarding transcript termination mechanisms across bacterial species.

While many studies are using transcript 3’-ends as signal for quantitative analysis, not all of these 3’-ends are created or captured in the same way. Typically, 3’-seq identifies signal from 3’-ends by ligating sequencing adapters to the 3’-ends of transcripts prior to fragmentation^13^. This approach captures signal from the 3’-ends of full-length and prematurely terminated transcripts, as well as transcript 3’-ends created by exonucleases, cleavage, damage, or other biological processes. This broad range of biological processes resulting in free 3’-ends introduces noise into 3’-seq data. Genomic coordinates with no known or likely biological function often display low levels of 3’-seq signal, suggesting, at face value, a small population of transcripts in the cell that terminate at such genomic positions. However, since not every genomic position can be the site of a preferred or regulated transcript boundary, analysis and interpretation of 3’-seq data typically relies heavily on the ability to differentiate true signal from biological or technical noise. This problem is heightened by the large range of expression levels for different transcripts, such that weak signals may be biologically relevant.

As the application of 3’-sequencing to address biological problems across diverse bacterial species continues to increase, so too does the number of 3’-seq analysis approaches. Many such analysis approaches are created “in-house” by labs and are designed to specifically analyze data from individual organisms for individual experiments and often make assumptions based on the known biology of the particular organism^2,3,5,7,16,17^. With each analysis tool using potentially different analytical or statistical frameworks and distinct biological assumptions, it is difficult to assess the biological relevance of results produced by two different methods, even when applied to the same data. This issue is exacerbated when attempting to compare results from two different studies (even on the same organism), as the differences in biological data derived from small variations in protocol or experimental design are compounded by inconsistencies in analysis approach.

To begin to address this issue, we previously developed PIPETS (<u>P</u>oisson Identification of <u>PE</u>aks from <u>T</u>erm-<u>S</u>eq data), a gene agnostic, statistically robust 3’-seq analysis method that is designed to analyze 3’-seq data of differing read depths from different organisms^18^. PIPETS employs a Poisson Distribution test to identify significant 3’-seq signal from surrounding noise while maintaining sensitivity by assessing coverage within a sliding window across the genome. Previously, using data from *Streptococcus pneumoniae* and *Bacillus subtilis*, we showed that PIPETS was able to identify significant 3’-seq signal across the genomes of both organisms, in gene contexts that are traditionally studied in the field (5’-UTRs and 3’-UTRs of annotated genes), as well as gene contexts that are not as often studied (gene coding regions, antisense to genes on the opposite strand, and intergenic regions).

With the goal of assessing the consistency of 3’-sequencing data derived from different studies conducted by different groups, we analyzed the raw 3’-seq data from three studies conducted with *Escherichia coli*^1–3^ using a common analysis strategy. During the course of our analysis, we uncovered large variations between the studies in both the reported results and the properties of the raw data. Overall, we find that most of the significant 3’-seq signal consistently identified by the three studies are from 3’-seq sites associated with tRNA synthesis rather than mRNAs, and that there are many instances where individual studies had significant 3’-seq signal that was comparably absent in other studies. Further, we identified substantial discrepancies in the 3’-seq read coverage distribution between the reference and complement strand data for many data sets. This is surprising given that the studies used similar extraction protocols, growth and experimental conditions, the same genome annotations and strains, and had similar sequencing read depths and qualities. Our findings constitute a call to either recognize the inconsistencies of such data in our interpretations, or to standardize both experimental and analysis approaches so that, as a field, we may move from a paradigm where each study exists in a vacuum, toward one where studies may together enable greater biological understanding of individual species and the differences between species.

## Materials and Methods

### Data Selection

All 3’-sequencing data was collected from sequence archives as .fastq files containing raw 3’-seq reads. The accession numbers for each data set used in this study are listed in Supplemental Table 1. We selected *E. coli* data sets that were sequenced from samples under WT experimental conditions or experimental conditions that were very similar to another of the three *E. coli* studies^1–3^. For the non-*E. coli* data sets, we selected 3’-seq data sets from a broader set of conditions to have enough samples for proper comparison.

### Alignment and Reference Genomes

All raw 3’-seq data sets were aligned using Burrows-Wheeler Alignment Tool^19^ (version 0.7.18-r1243) with default parameters. All data sets were aligned using the reference genomes provided in their respective studies. For the three *E. coli* studies we used the *E. coli* MG1655 (NC_000913.3) reference genome. For the other studies, we used the *B. subtilis* 168 Chromosome 1 (AL009126.3) reference genome for the Dar^13^ data, the *B. burgdorferi* B31 (GCA_000008685.2) reference genome, the *P. aeruginosa* UCBPP-PA14 (NC_008463.1) reference genome, and the *S. pneumoniae* TIGR4 (NC_003028.3) reference genome. Following alignment with BWA, all .sam files were converted to .bam files using bedtools^20^ (v2.30.0) view with default parameters, and all .bam files were converted to .bed files using bedtools bamtobed with default parameters. The .bed files were then used as input files for PIPETS analysis.

### Analyzing Selected Data Sets with PIPETS

PIPETS (version 1.4.6) was run with different values of the parameters “threshAdjust” and “highOutlierTrim” to identify the level of analytical strictness that best suited each data set. Both parameters are used to calculate the minimum number of 3’-seq reads (read coverage cut-off) required for a genomic coordinate to be tested for significance by PIPETS based on the properties of an individual dataset. The value of threshAdjust determines the percentage of genomic coordinates whose read coverage values (starting with highest coverage) are averaged to determine the minimum read coverage cut-off value. The value of highOutlierTrim determines the percentage of the highest expressing genomic coordinates selected via threshAdjust that are removed from calculation of the read coverage cut-off to reduce the influence of high expressing outliers. Generally higher values of threshAdjust increase sensitivity.

Parameter values were chosen to maximize the number of significant peaks identified without lowering the minimum read coverage cutoff to values that were below our confidence threshold for having potential biological significance. The threshAdjust and highOutlierTrim values used for each data set and replicate can be found in Supplemental Table 2. Although PIPETS can analyze the reference and complement strand of a data set separately, we chose to analyze both with the same criteria to capture any strand specific differences in the data. For all PIPETS analysis done here, we set readScoreMinimum = 30 (read quality) and did not change any parameters other than threshAdjust and highOutlierTrim from their default values.

### Combining Results from PIPETS

We first took the significant *E. coli* PIPETS results of the replicate data sets from each condition and identified consistently present 3’ sites. For studies with 3 replicates per condition, a significant 3’ site needed to be found in 2 of the 3 replicates (an overlap distance of +/-10 bp was used). In studies with 2 replicates per condition, a significant 3’ site needed to be found in both replicates (overlap distance of +/- 10 bp was used). This list of 3’ sites was then used as the study-specific results for this work (Supplemental Table 3).

For the *B. subtilis*, *P. aeruginosa*, *B. burgdorferi,* and *S. pneumoniae* data we followed the same procedure for combining significant identified 3’-sites as above. As some of these data had multiple conditions other than WT, we chose to additionally filter the 3’-sites that were found in sufficient replicates by removing any that were not found in 66% or more of the conditions for that study (e.g. in the instance of the Dar data which has 9 conditions, a 3’-site must be identified in a sufficient number of replicates for 6 or more conditions).

### Classifying Results under genic contexts

Given the inherent differences between organisms, we wanted to create data informed classification systems for important genomic contexts in the different organisms. We used the genomic coordinates of the significant PIPETS results to establish metrics for the average distance of the 5’UTR and 3’UTR from the nearest gene coding region. For each study, we calculated the arithmetic mean distance of the significant PIPETS 3’-sites (only those that were present in 66% or more replicates, see above, and were within 1000 bp of a start or stop site) from the nearest gene coding region start sites (5’-UTR) (e.g. Adams Strand 5’UTR Mean Distance) and stop sites (3’-UTR) (e.g. Adams Strand 3’UTR Mean Distance). This was performed independently for the reference and complement strands using strand specific 3’-sites. We then calculated the 5’-UTR and 3’-UTR distance metrics for each strand by taking the geometric means of the previously determined values from each study (Equation 1,2).

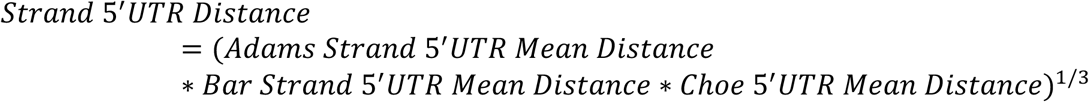

**Equation 1: 5’-UTR distance metric calculation using geometric mean.** “Strand” refers to either the reference or complement strand as this equation was calculated independently using the arithmetic mean distance values from each study in a strand specific manner.

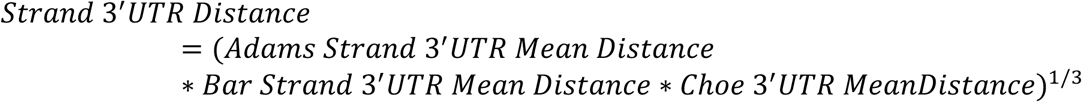

**Equation 2: 3’-UTR distance metric calculation using geometric mean.** “Strand” refers to either the reference or complement strand as this equation was calculated independently using the arithmetic mean distance values from each study in a strand specific manner.

The PIPETS results were then assigned as being Genic if they were inside of any coding region in the annotation, 5’UTR if they were within the established distance from the start of a coding up to 10bp into a gene, 3’UTR if they were within the established distance from the stop of a coding region up to 10bp into a gene, Antisense if they were within the coding region of a gene on the opposite strand and Intergenic if they were not categorized as Genic, 5’UTR, and 3’-UTR. Sites may be classified in multiple contexts (e.g. 3’-UTR and genic in the downstream gene or Antisense and Intergenic).

### Outside Methods to Further Classify PIPETS Results

We used three terminator prediction tools to further study the 3’ sites identified by PIPETS. For Rho-dependent terminator prediction, RhoTermPredict^21^ (Version 3.4.0) was run on the *E. coli* MG1655 (NC_000913.3) reference genome with default parameters. For Rho-independent terminator prediction, we used RNIE^22^ (Version 0.01) and TransTermHP^23^ (Version 2.07). RNIE was run on the *E. coli* MG1655 (NC_000913.3) reference genome with default parameters. TransTermHP required a .ptt file for analysis along with the *E. coli* MG1655 reference genome for analysis. We used gb2ptt (https://github.com/sgivan/gb2ptt) to convert the *E. coli* MG1655 GenBank File (U00096.3) to a .ptt file as the required input for TransTermHP which was run with default parameters. We considered all results for TransTermHP and RNIE, but we only used the RhoTermPredict results which had a score_sum value of 160 or greater for comparative analysis. For the results of all three tools, we checked if any PIPETS significant 3’ site was found within 150 bp downstream (relative to strand) of the predicted terminator. If so, we considered those to be overlapping results.

## Results

### Individual studies report varying sets of significant 3’ ends

In order to provide a baseline expectation for how the results of 3’-sequencing studies performed by different groups under similar conditions would overlap for *E. coli*, we compared the published lists of termination sites from 3 studies. For this work, we chose three *E. coli* studies, Adams et al.^1^ Bar et al.^2^ and Choe et al.^3^, because each had multiple 3’-seq replicates with read depths greater than 10^7^ reads. Additionally, despite the presence of other studies which use data labelled as 3’-seq/Term-seq, these three studies were the only relevant works to directly perform 3’-end sequencing on the same organism as opposed to deriving 3’-ends from RNA-seq or other techniques^24–28^. Since each of these studies also performed other types of sequencing, we selected only 3’-seq/Term-seq/3’-end data sets from wild-type strains, which resulted in 6 Adams, 6 Bar, and 3 Choe 3’-seq data sets. The conditions for these samples were all very similar; the Adams and Bar studies used LB media as well as a minimal media condition, while the Choe data did not specify the medium conditions under which their Term-seq data sets were grown. All three of the *E. coli* studies cultured the MG1655 strain to mid- or late-log and used the same reference genome (NC_000913.3) for their downstream analysis.

In assessing the reported sites, we selected, to the best of our ability, only the published results associated with the conditions listed above from each of the studies. Given that each study used the same reference annotation, similar preparation and sequencing protocols, and similar growth conditions, we concluded that the generation of the 3’-seq data from each study should result in comparably similar 3’-seq data and results. When comparing the reported sites, we examined the data in a strand-specific manner and considered sites reported within +/- 5 bp to be overlapping to provide a generous interpretation of overlap.

The published results of the *E. coli* studies had more than 200 sites that were found in all three studies for both the reference and complement strands (Figure 1A & 1B). While there was still overlap in a pairwise manner between the Adams, Bar, and Choe published results, a majority of the published results were unique to individual studies. The Bar study in particular had more than 700 unique results per strand, which is almost comparable to the total number of published results for either of the other two studies.

**Figure 1:**
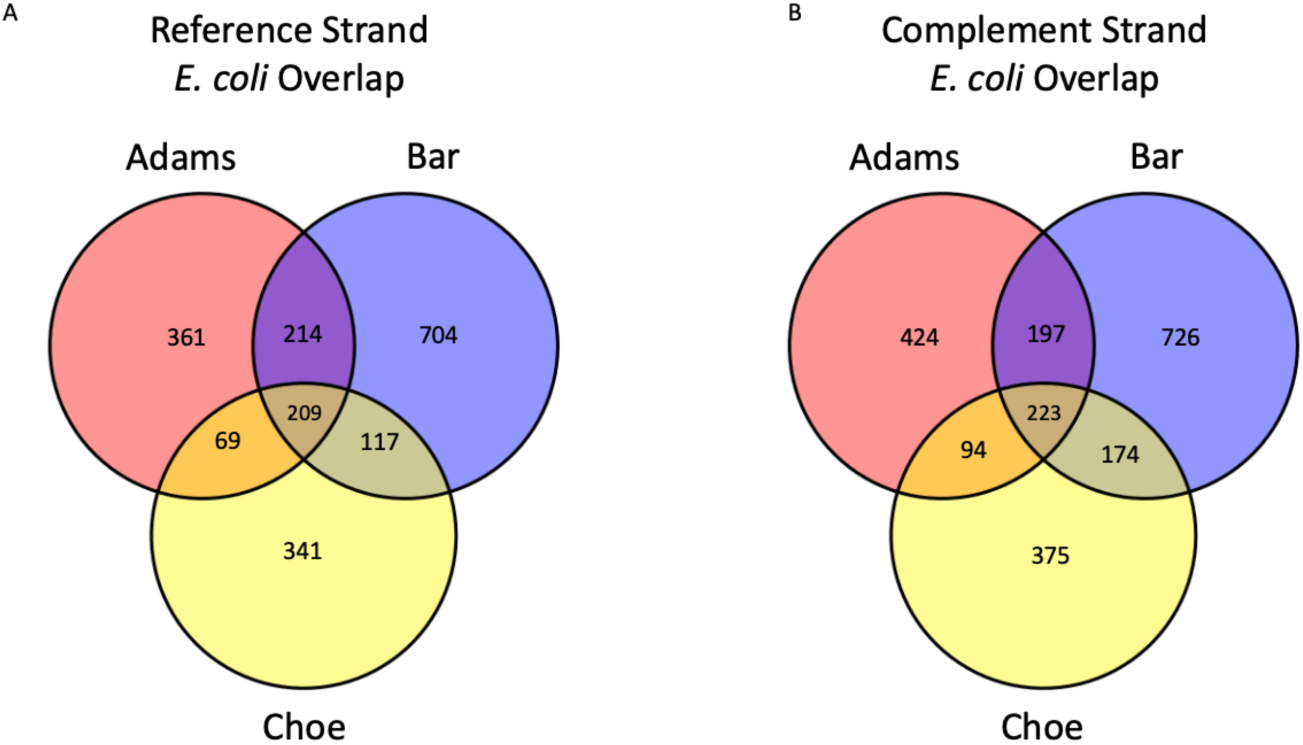
Published results of *E. coli* studies show some overlap, but each study still has many more unique results than shared. A) The reference strand published results of the Adams, Bar, and Choe studies had 209 3’-seq sites that were present in all three studies along with varying levels of pairwise shared results. However, each study had at least 300 results that were unique to that study, with the Bar data having more than 700 unique 3’ sites. B) The complement strand published results of the Adams, Bar, and Choe studies showed the same trend as the reference strand published results, with even more study-specific results for each of the three studies. List of 3’termination site coordinates reported in Supplemental Table 4.

This comparison illustrates the challenges associated with combining multiple studies to gain greater biological insight. While there are some variations between the studies, the similarities between the growth conditions and reported experimental protocols used for 3’-sequencing do not suggest that these studies should yield such disparate results. Thus, we were surprised to see such a high number of study-specific results in comparison to the overlapping 3’-seq sites. This comparison, at face value, suggests that there are ∼1400 unique, biologically relevant 3’-seq sites on both strands of the *E. coli* genome that were generated under very similar experimental conditions and that each study captures ∼50-85% of these. As this seems unlikely, we cannot determine if these discrepancies are the result of differences in the raw 3’-seq data due to: minor changes in experimental or sequencing protocols, the 3’-seq analysis methodology chosen, or the decisions surrounding the interpretation of the results.

### PIPETS Parameterization Data Reveals Broad Diversity of 3’-seq Data Characteristics

In order to remove as many confounding factors as possible from our comparison, we collected the raw 3’-seq data of each of the studies and sought to analyze and compare the results in an equivalent manner. To do this, we employed a 3’-seq analysis package previously developed in our lab called PIPETS^18^. PIPETS (<u>P</u>oisson Identification of <u>PE</u>aks from <u>T</u>erm-<u>S</u>eq data) is a genome annotation agnostic, statistically robust 3’-seq analysis package available on Bioconductor. We previously demonstrated that, of the parameters PIPETS uses to adjust sensitivity, the parameter “threshAdjust” is the most impactful for changing the set of significant sites returned by PIPETS. threshAdjust is used to calculate a minimum read coverage cutoff that must be exceeded in order for any genomic coordinate to undergo a Poisson Distribution test for significance. Many studies use an arbitrary minimum read cut-off in determining significance of a particular site; threshAdjust allows this cut-off to be defined by the read-depth of an individual dataset. This dynamic cut-off is essential for maintaining sensitivity and ignoring low-confidence 3’-seq signal when analyzing data-sets with different read depths (Figure 2A). While the read depth and read coverage distribution of each 3’-seq file is unique, we do consistently observe that increasing the value of threshAdjust (making PIPETS analysis less strict and potentially more sensitive) increases the number of identified significant 3’-seq sites, while decreasing the read coverage cut-off for 3’-seq data (Figure 2B & 2C). We noted that, for the individual replicates depicted, the Adams data had fewer significant 3’ sites detected than either the Bar or Choe data which was exaggerated at higher levels of threshAdjust (Figure 2B). This corresponded with higher read coverage cut-off values for the Adams data compared to both the Bar and Choe data (Figure 2C).

**Figure 2:**
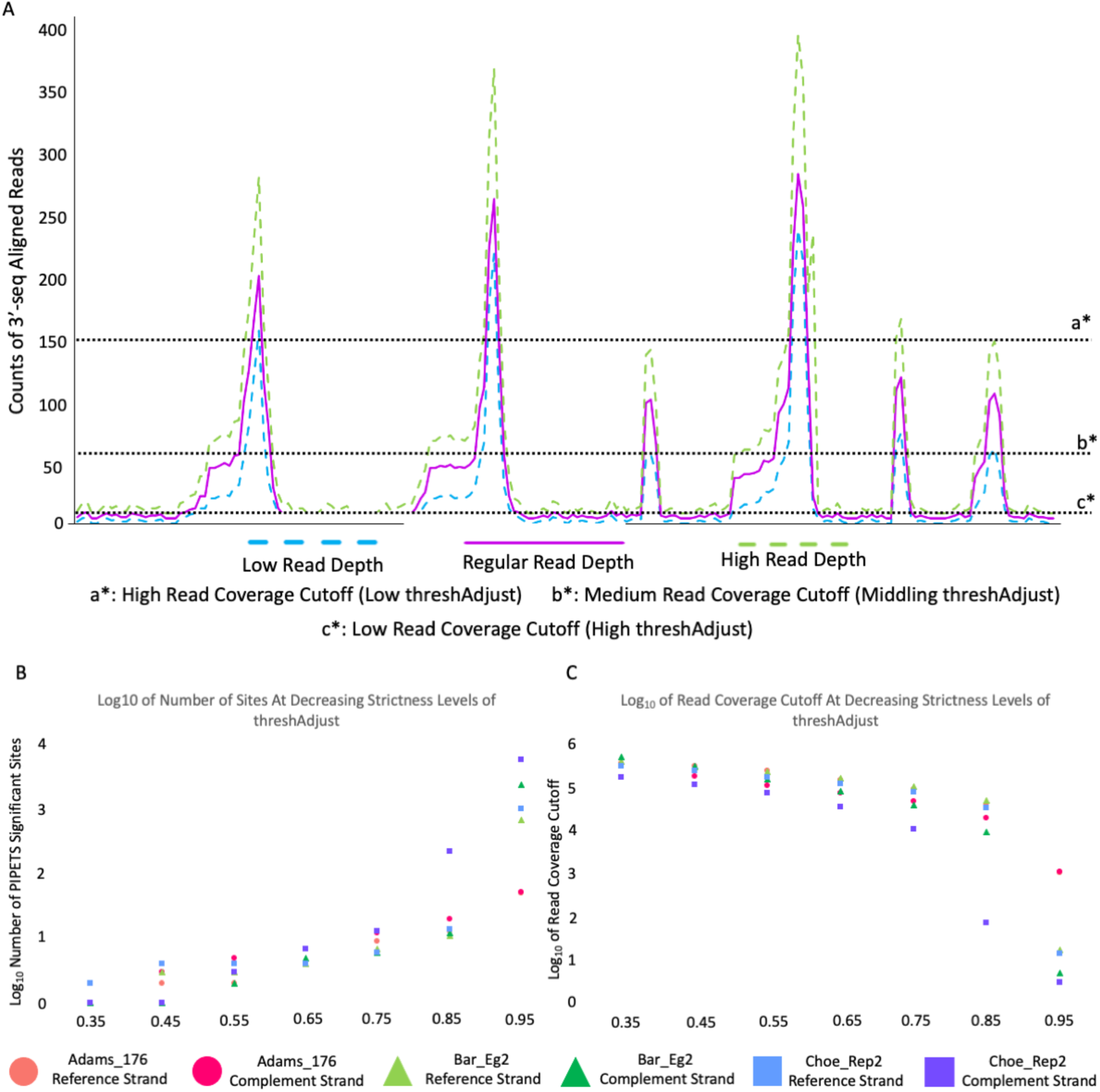
Differing levels of strictness of PIPETS analysis are better suited to analyze different data sets. A) Artificial demonstration 3’-seq data showing terminal position read counts for three possible read depth scenarios: low, medium, and high. Different read coverage cutoff values of PIPETS, indicated by the dotted lines, are better suited for different data sets with different read depth and noise levels. B) One representative replicate was chosen from each *E. coli* data set for demonstrative graphing. As PIPETS is made less strict (threshAdjust increases), more significant sites are identified. We note that these replicates of Bar and Choe data have many more 3’-seq sites identified at higher levels of threshAdjust. C) The read coverage cutoff required to be identified as significant drops as PIPETS becomes less strict (threshAdjust increases). Following the trend from the Figure 2B, the Bar and Choe read coverage cutoff is much lower than the that of the Adams data replicate.

To determine if the trends seen for individual replicates for each study were consistent, we analyzed each of the studies with PIPETS at three different values of threshAdjust (0.5, 0.75, 0.95) to identify the 3’-seq profile of each study. The reference and complement strand parameterization results both showed the same consistent trend: as threshAdjust increases, more significant 3’-sites are identified (Figure 3A & 3C) and the read coverage cut-off of PIPETS decreases (Figure 3B & 3D). As seen with the individual replicates in Figure 2B/C, the Adams data has consistently fewer significant 3’-sites than both the Bar and Choe data at high values of threshAdjust (Figure 3A & 3D), which corresponded with higher read cut-off values (Figure 3B & 3D). This difference was substantial; on the reference strand at threshAdjust = 0.95, the Bar and Choe data each had 10-30 times more significant 3’-sites than the Adams data, and both had read coverage cut-off values that were one-to-two orders of magnitude lower than the Adams data (Supplemental Table 5). The complement strand parameterization data was even more dramatic at high values of threshAdjust; the complement strand Bar data had up to 100 times more significant 3’-sites than the Adams data with read coverage cut-off values that were two-to-three orders of magnitude smaller. The complement strand Choe data was the most extreme, with one replicate having more than 8000 identified significant 3’-sites and a read coverage cut-off of 2 reads. The large discrepancy between the Adams data and the Bar and Choe data strongly indicates an underlying difference in distribution across the genome of the 3’-seq data of these studies: the Adams data has fewer 3’-sites that account for a greater proportion of the total 3’-seq read coverage, while the Bar and Choe data have a wider distribution of 3’-seq reads across all genomic coordinates.

**Figure 3:**
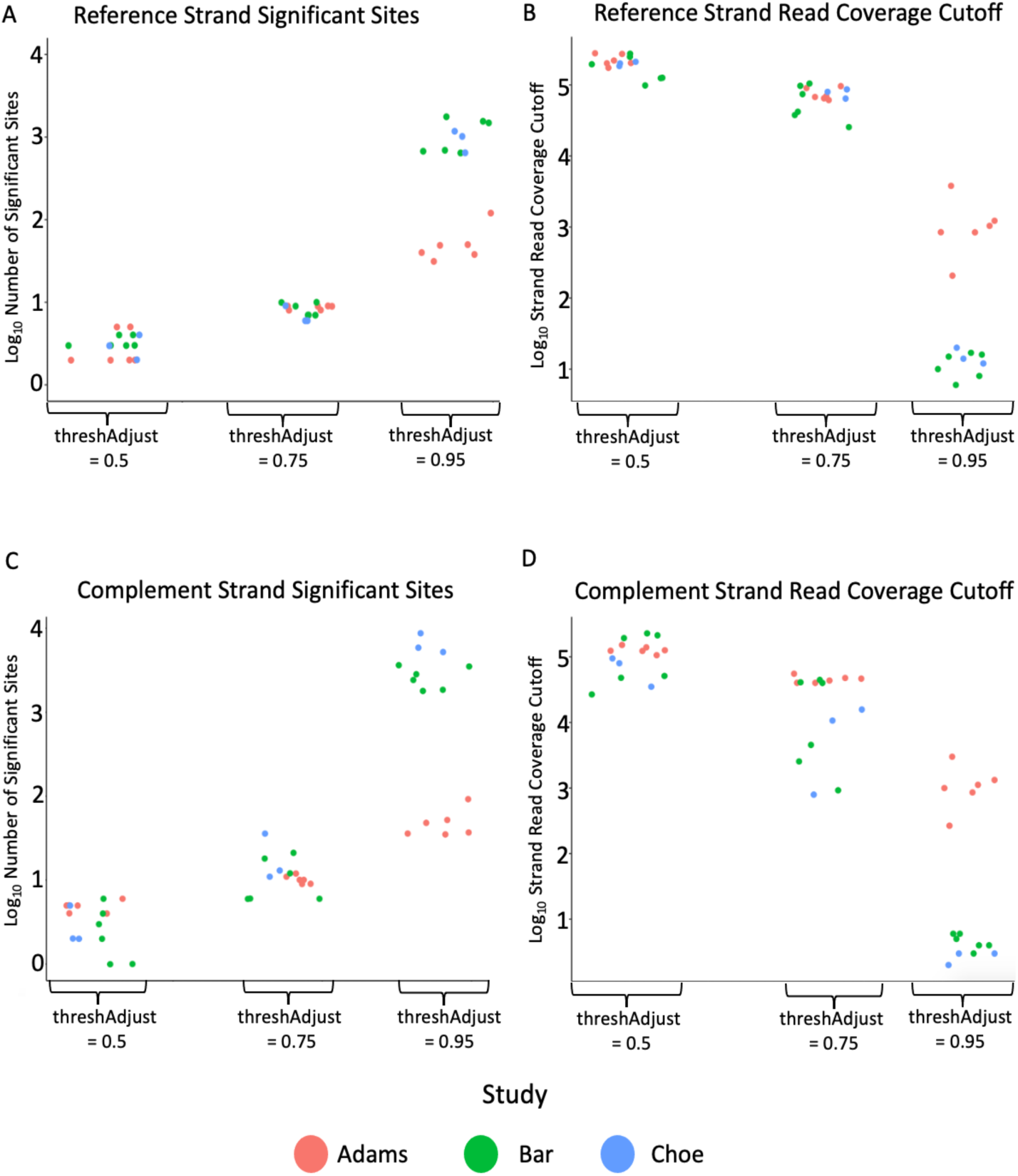
Reference and complement strand mapping *E. coli* 3’-seq data show different patterns at varying levels of threshAdjust, which is more pronounced on the complement strand. A) PIPETS was run at three different values of threshAdjust for all Adams, Bar, and Choe data sets. highOutlierTrim set to 0.01 these analyses. The log_10_ number of identified reference strand 3’ sites was consistent for threshAdjust = 0.5 and = 0.75, but the Adams data had fewer 3’ sites at threshAdjust = 0.95. B) The log_10_ read coverage cut-off value for the reference strand data was taken for the same data sets at the same values of threshAdjust. A similar pattern to the number of 3’ sites was noted with the Bar and Choe data having dramatically lower log_10_ read coverage cut-offs at threshAdjust = 0.95. C) The log_10_ number of complement strand 3’ sites had a similar trend to the number of reference strand 3’ sites, however the difference between the number of Adams 3’ sites and Bar and Choe 3’ sites was even greater for the complement strand results. D) The complement strand log_10_ read coverage cut-off values were even more different than the reference strand results, with the Bar and Choe results having much lower read coverage cut-off values than the Adams data, which was noticeable even at threshAdjust = 0.75.

The separation of the Adams data from the Bar and Choe data suggests that these two clusters are either capturing entirely different 3’-seq signal or there are mitigating factors that are affecting the population of 3’-seq reads that are being sequenced. The Bar and Choe results have orders of magnitude more identified 3’-sites than the Adams results, a difference that is amplified for the complement strand results (Supplemental Table 5). It is impossible to determine which, if any, of these three data sets is best capturing the true 3’-seq profile of *E. coli* under these similar conditions. Given these irregularities, we sought to identify potential candidate causes that could have contributed to the pervasive differences in 3’-seq profiles. We compared the distribution of mapping-quality passing 3’-seq reads across the *E. coli* genome for each sample, and noted that while the Adams samples generally had more genomic coordinates with 1000 or more mapped 3’-seq reads than either the Bar or Choe data, though the difference was not so great to wholly explain the changes in parameterization data (Supplemental Table 6). While all of the Bar and Choe raw 3’-seq data had more reads mapping to the reference strand than the complement strand, the difference was not sufficient to fully explain the dramatic increase in identified sites and decrease in read coverage cut-off at higher levels of threshAdjust (Supplemental Table 7).

This complement strand skew is surprising given that the distribution of *E. coli* gene coding regions is roughly even between the reference and complement strands^29^, suggesting that the discrepancy noted in these studies might not be the result of biological processes. Given that the three experiments were conducted with very similar conditions, if the data accurately reflect the frequencies of biologically relevant 3’-ends, we would expect to see all three studies clustering together at every value of threshAdjust. While the separation becomes most apparent at threshAdjust = 0.95, we see the clustering break apart at threshAdjust = 0.75, suggesting that the differences in the characteristics of these data sets is not exclusively found in the presence or absence of low read coverage positions (Figure 3C & 3D). Despite the apparent differences, there are some unifying factors among the data of the three studies: there are a small number of very high read coverage genomic positions that dominate the *E. coli* 3’-seq read coverage distribution and the influence of low read coverage noise positions is low, with no more than 0.02% of genomic coordinates having an identified significant 3’-site in the even the highest 3’-site count among the Choe data (Supplemental Table 5).

### E. coli Significant PIPETS Results Are Few in Number and Have Very Little Overlap

The heterogeneity of the 3’-seq signal for the *E. coli* data sets made it difficult to identify if the differences between the PIPETS results for each study were caused by biologically relevant changes or simply inconsistencies in 3’-seq read depth and quality. We sought to better understand how the 3’-seq results of these studies differed and if there were notable patterns that could be identified. We ran PIPETS on each data set with parameter values that that best captured the attributes of the data while also identifying significant 3’-sites with read coverage values that we believed could be biologically relevant (Supplemental Table 2). We took all significant sites that were identified in a majority of replicates for each *E. coli* study individually, and categorized these sites as 5’-UTR, 3’-UTR, Genic, Intergenic, and Antisense (see methods). We saw that most significant sites had multiple potential genetic contexts (e.g. Genic and 5’-UTR of the following gene), and that the reference and complement strand distribution of genetic contexts differed, with the complement strand having a higher population of sites with 5’-UTR or 3’-UTR genetic contexts (Supplemental Figure 1). We next compared these sites amongst each other to identify any consistent overlap between the studies.

We found little overlap between the PIPETS results of the three *E. coli* data sets, with very few sites being present in a sufficient number of replicates for any study, with the exception of the Choe complement strand results (Figure 4A). The low number of significant sites was robust to changes to reasonable threshAdjust and highOutlierTrim, particularly for the reference strand, suggesting that for each of these data sets there were a handful of genomic positions that accounted for a majority of the 3’-seq signal.

**Figure 4:**
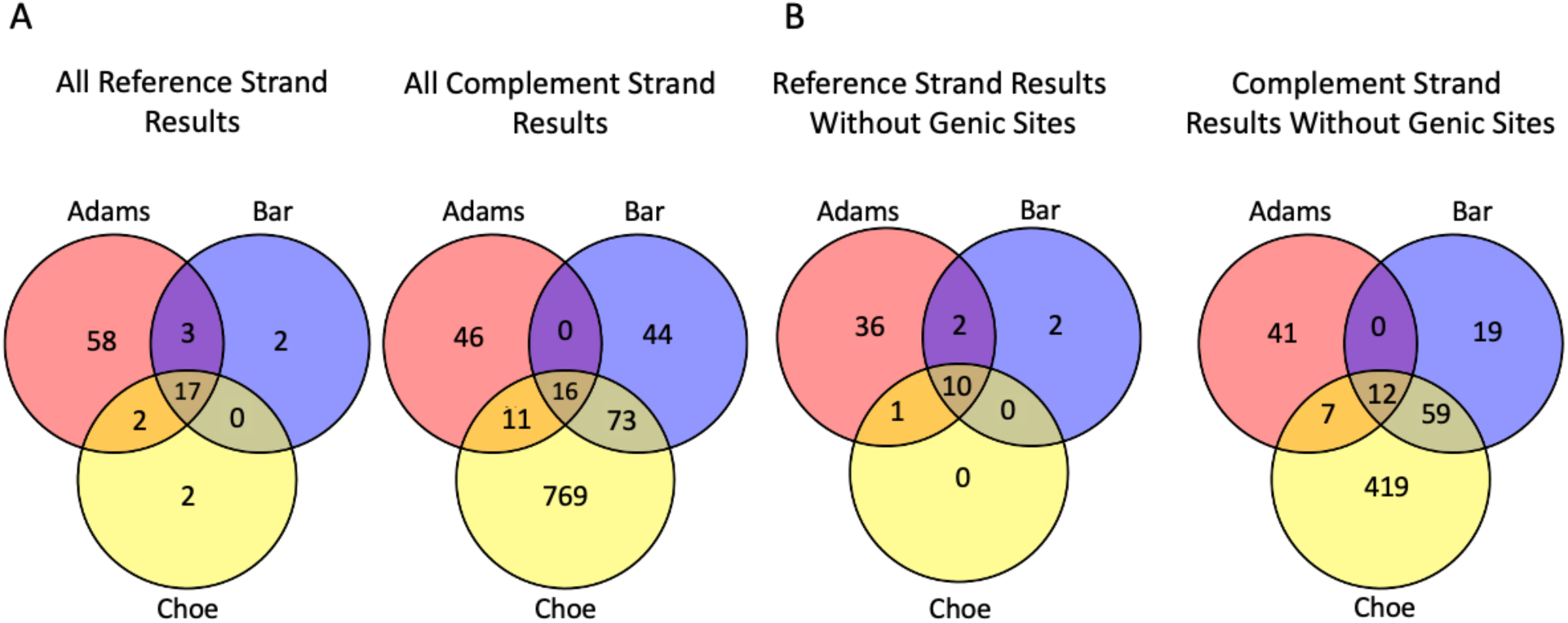
PIPETS results from *E. coli* data sets show very little overlap, which is not substantially changed by removing genic signal. A) We compared the PIPETS results that were present in 66% or more of replicates for each study and found very little overlap between any of the *E. coli* data sets. The largest outlier in the analysis was the Choe complement strand data, which had 769 sites that were unique to that study. B) We removed all PIPETS sites that were found inside of gene coding regions, regardless of their other potential genomic functions. We found no large shift in the trends of the overlap, and although the Choe complement strand data was altered the most, it was still a dramatic outlier in terms of unique 3’-seq results.

In order to capture as many 3’-sites as possible for overlap comparison, we elected to increase highOutlierTrim to values much higher (values from 0.1-0.5) than the default (0.01) instead of further adjusting threshAdjust. This step removes more sites of high coverage from our calculation of the read-depth cutoff (they are still tested for significance) and thus decreases the read coverage cut-off and increases the number of identified 3’-sites by reducing the impact of high read coverage positions on the analysis. However, the resulting additional genomic sites did not overlap between studies, despite the large number of 3’-sites identified in the Choe complement data.

To identify the cause of the high number of Choe complement strand sites, we removed any significant peak that was present inside of the gene coding region of any gene, regardless of if it had other potential genomic roles. This practice has been employed in several studies focused solely on 3’-seq signal present in the 5’ or 3’-UTR of annotated genes. We compared the overlap of the three studies without any genic sites and saw an identical trend for both the reference and complement strands (Figure 4B). While the largest reduction in sites was for the Choe complement strand results, there are still 419 sites that are not present in the results of either the Adams or the Bar data.

For each study, we compared the number of unique 3’-sites that were present before and after the replicate presence threshold (without removing Genic sites). In order to pass the replicate presence thresholding, a unique 3’-site must be identified as significant in 66% of the replicates for a study (i.e. 4 of 6 Adams replicates, 4 of 6 Bar replicates, or 2 of 3 Choe replicates). The Adams data had consistent counts of unique 3’-sites between the reference and complement strand results, and both strands lost comparable amounts of 3’-sites after the replicate threshold (Table 1). The Bar and Choe data both had considerably more complement strand 3’-sites than reference strand 3’-sites before the replicate threshold. However, while 869 of the 1344 Choe complement strand 3’-sites passed the replicate threshold, only 133 of the 1082 Bar complement strand 3’-sites passed. Although the Choe data had three replicates compared to the six Bar data replicates, the large disparity in replicate passing complement strand 3’-sites (∼12% of the Bar complement strand sites vs. ∼65% of the Choe complement strand sites) suggests broader inconsistencies in 3’-seq read coverage distribution between replicates from the same study. Even if the strictness of the replicate threshold is reduced such that a 3’-site must be detected in two total Adams or Bar replicates instead of four, there is still little overlap between all studies, although the overlap between the Bar and Choe studies does increase (Table 1) (Supplemental Figure 2).

**Table 1:** Unique 3’-sites present before and after replicate presence threshold for each study. Values in the denominators denote the total number of strand specific unique 3’-sites across all replicates of each study. The Choe data had 3 replicates, so no 33% threshold was used, as only one replicate did not provide sufficient confidence.

| Study | Reference 3'-sites<br>Passing 33%<br>Replicate<br>Threshold | Complement 3'-sites<br>Passing 33%<br>Replicate Threshold | Reference 3'-sites<br>Passing 66%<br>Replicate<br>Threshold | Complement 3'-sites<br>Passing 66%<br>Replicate Threshold |
| --- | --- | --- | --- | --- |
| Adams | 127/182 = 69.8% | 101/143 = 70.6% | 80/182 = 44% | 73/143 = 51% |
| Bar | 35/50 = 70% | 634/1082 = 58.6% | 22/50 = 44% | 133/1082 = 12.3% |
| Choe |  |  | 21/24 = 87.5% | 869/1344 = 64.7% |

With very few sites passing the replicate presence threshold relative to the number of total 3’-sites, it is difficult to determine if this overlap is truly representative of the similarity between the data from these three *E. coli* studies. We next identified what significant 3’-seq sites were found in all three studies on both strands. Using RNAcentral^30^, we took the 150 bp region upstream (relative to strand) of the PIPETS peak and tested if those sequences aligned to any known regulatory elements. We found that the *E. coli* results that were present in all three studies predominantly matched with tRNA functions (RF00005) and some matched to signal recognition particle functions (RF00169). This suggests that the significant sites found between all three *E. coli* studies come from signal from high abundance transcripts (since tRNA’s are not depleted in most 3’-seq protocols) and not necessarily from signal from transcripts of protein coding genes.

To investigate the potential consequences of these large 3’-seq read coverage disparities, we examined the 3’-seq read pileup of the aligned 3’-seq data from one of each of the *E. coli* studies using Integrated Genome Viewer (IGV)^31^. We noted large differences in both read coverage magnitude and read coverage distribution for given regions of the genome, but these changes between studies were not always consistent. In the region spanning from ∼1958kb - ∼1974kb on the *E. coli* MG1655 genome, we saw a single, standard termination peak at ∼1958.5kb for the Choe data, with little to no noise in the rest of the region (Figure 5A). The Bar data had ∼20% of the total magnitude of reads in this region, and featured a very small peak at that same location. It instead had a large amount of noise and other signal in genomic coordinates that were not seen at all in the Choe data. The Adams data had almost no reads mapping to this stretch of the genome. When we compare the read coverage cutoffs generated for these individual files with their IGV read pileups, we can see that the Choe cutoff of 16 3’-seq reads, which was much lower than the other two, would have only identified a few total sites for its own data, given the sparse noise signal. It is expected that the raw complement strand reads, which are displayed in Figures 5 and 6, would be associated with reference strand transcription events. None of the three data sets had 3’-seq read pileup peaks that were proximal to any reference strand genes (marked in blue arrows), instead almost all of the 3’-seq read pileups, both peaks and noise, were associated with complement strand gene coding regions. This suggests that the complement strand 3’-seq reads in this region have antisense functionality (Figure 6A).

**Figure 5:**
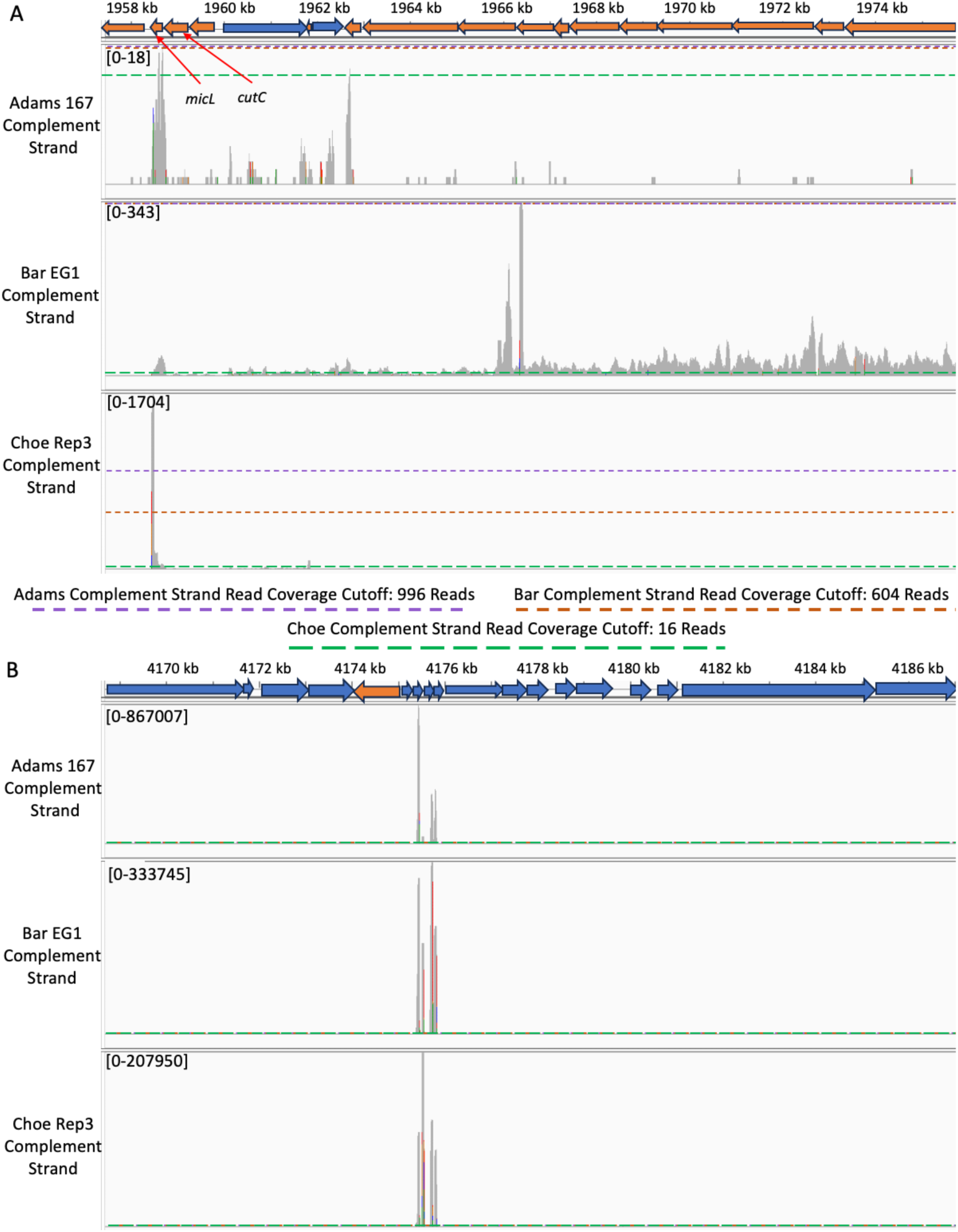
IGV read pileup graphs for example data sets from the Adams, Bar, and Choe *E. coli* studies show discrepancies in read coverage and expression patterns. A) IGV read pileup graphs (read lengths of studies were 37, 77, and 49 respectively) for one representative complement strand data set of each of the Adams, Bar, and Choe studies. The complement strand reads correspond to signal generated from reference strand transcription due to the 3’-seq protocol, so 3’-seq read pileup peaks should align with blue arrows if they are cis-generated. The read coverage cutoffs were generated with threshAdjust values of 0.95, 0.9, and 0.85 respectively, and a highOutlierTrim value of 0.01 for all replicates. Blue arrows indicate reference strand gene coding regions and orange arrows indicate complement strand gene coding regions. The Choe data is the only one to feature a prominent peak near the end of the *cutC* and *micL* genes (∼1958.5 kb) on the complement strand. The Adams and Bar data have substantially lower total read coverage in this region, but not a comparably lower total file read depth, and the expression patterns of all three data sets are different and distinct. The Bar data had large levels of antisense noise across the entire region, with no substantial peak aligning to any reference strand gene coding regions. B) IGV read pileup graphs show inconsistent read coverage and expression patterns for four tRNA genes (*thrU*, *tyrU*, *glyT*, and *thrT*) at ∼4175.5 kb. All cutoffs are present at the foot of the graphs as they are well below the relative read coverage magnitude of the region. The Adams data has ∼2.5-4 times higher read coverage magnitude in this region, but its relative expression profile is different than either the Bar or the Choe data.

**Figure 6:**
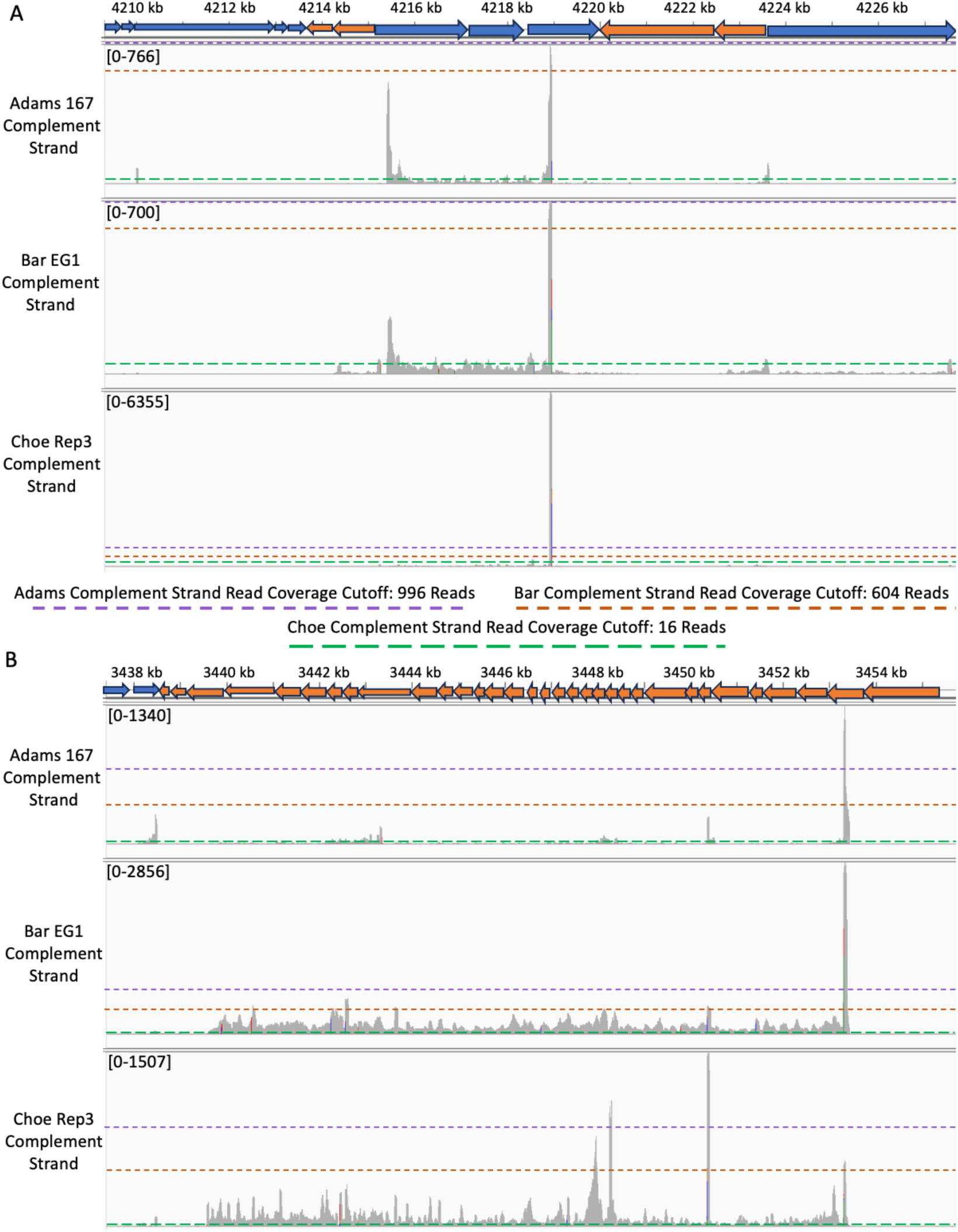
IGV read pileup graphs reveal inconsistencies in noise distribution and differences in potential primary termination sites for each study. A) IGV read pileup graphs (read lengths of studies were 37, 77, and 49 respectively) for one representative data set of each of the Adams, Bar, and Choe studies. The complement strand reads correspond to signal generated from reference strand transcription due to the 3’-seq protocol, so 3’-seq read pileup peaks should align with blue arrows if they are cis-generated. The read coverage cutoffs were generated with threshAdjust values of 0.95, 0.9, and 0.85 respectively, and a highOutlierTrim value of 0.01 for all replicates. Blue arrows indicate reference strand gene coding regions and orange arrows indicate complement strand gene coding regions. Although all three data sets have a primary 3’-seq read pileup peak at ∼4219 kb, only the Adams and Bar data have any noise signal in the preceding region, while the Choe data has almost 10 times the total reads mapping to the location of the peak. B) The Bar and Choe data have a large stretch of noisy antisense signal across almost the entire region, but only the Adams and Bar studies have a primary 3’-seq read pileup peak at ∼ 3453kb. The Choe data has three dominant peaks, only one of which (∼3450 kb) is present in any meaningful capacity in the other two studies.

We also identified instances where the differences in 3’-seq read coverage were affecting regions with very high read coverage positions and little noise. In the region spanning from ∼4170kb – 4186kb on the same annotation, we saw several consecutive 3’-seq read pileup sites that aligned between the three data sets that were near the *thrU*, *tyrU*, *glyT*, and *thrT* tRNA genes (Figure 5B). However, we see that the magnitude of the read coverage for each of the studies is quite different, with the Adams data having a read coverage that was ∼3-4 times greater than the other two studies. Additionally, we noted that while each of the studies has roughly the same number of sites in the same region, the relative magnitude of each of the sites is noticeably different. Because the read coverage of these sites is so much higher than any of the read coverage cutoffs of the studies, there would be likely be no difference in the number of sites identified if different cutoffs were used for different studies.

We identified instances where despite similar significant peaks called by PIPETS, the captured 3’-seq read pileup distributions in the raw data diverged so heavily that qualitative analysis suggests substantial differences 3’-seq termination patterning. In the region spanning ∼4210 kb – 4226 kb, the Adams and Bar studies have 3’-seq signal read coverage patterns that were indicative of a classical gene termination pattern ending at ∼4219 kb (Figure 6A). There is some additional signal apparent within the preceding coding region in these datasets including a potential peak at ∼4215kb, which falls beneath the minimum read cut-off for both these datasets. However, the primary termination site for the Choe data (at ∼4219 kb) has nearly 10 times the 3’-seq read pileup of the Adams or Bar data, and the 3’-seq signal present in genic region, including the read pileup peak at ∼4215 kb is not present.

Finally, we saw cases where the 3’-seq read pileup was so different that no two studies presented the same 3’-seq signature. In the region spanning ∼3438 kb – 3454 kb, we note dramatically different distributions of 3’-seq read pileup for all three data studies (Figure 6B). The Bar and Choe data have a long stretch of noisy 3’-seq read pileups for a majority of the region, but the Adams data only has five regions with any 3’-seq read pileup. While the predominant signal for the Adams and Bar data is a read pileup peak at ∼3453 kb, the Choe data instead has three 3’-seq sites that are either lowly expressed or entirely absent in the other studies. Almost all of the 3’-seq read pileup coverage in this region for the three studies appears to be antisense signal, as the read pileup peaks are proximal to complement strand coding regions and not reference strand coding regions (Figure 6B). Although the Bar data has roughly double the read pileup magnitude in this region, it is surprising that the three studies differ not only in noisy read coverage distribution, but they also seem to have different primary termination sites despite similar experimental conditions.

### Additional 3’-seq Results from Different Model Organisms Do Not Disrupt E. coli Results Clustering Pattern

In order to test if the magnitude of difference between the three *E. coli* studies was severe when compared to data from other organisms, we analyzed four more 3’-seq data sets from four different model organisms: *Bacillus subtilis*^13^, *Pseudomonas aeruginosa*^7^, *Borrelia burgdorferi* ^5^, and *Streptococcus pneumoniae*^8^. The differences in 3’-seq data characteristics were present at all values of threshAdjust, with the Warrier^8^ and Konikkat^7^ data having dramatically more identified 3’-sites than any other study (Figure 7A & 7C) (Supplemental Table 8). Even with the addition of other data sets, the Adams data and the Bar and Choe data still fell into different clusters. The Adams and Petroni^5^ data clustered separately from all of the other data sets, and were the least variable between the reference and complement strands. Interestingly, we see that the Dar, Konikkat, and Warrier data all have a complement strand bias, with more total 3’-sites (Figure 7A & 7C) and lower read coverage cut-offs than the reference strand data (Figure 7B & 7D). The complement strand bias noted in the additional data sets has a comparable magnitude to the bias seen in the Bar and Choe data, suggesting that factors influencing this skew are pervasive across organisms or across experimental procedures of different studies. These comparisons further complicate our ability to determine which, if any, of the *E. coli* studies analyzed here represent an “accurate” set of 3’-seq characteristics for the organism under the condition assessed. The addition of four different organisms, two of which are Gram positive, contextualized, but did not explain, the discrepancies found between the three *E. coli* studies: the Adams data was most similar to *B. burgdorferi* 3’-seq data, and the Bar and Choe data were most similar to 3’-seq data from two Gram positive organisms.

**Figure 7:**
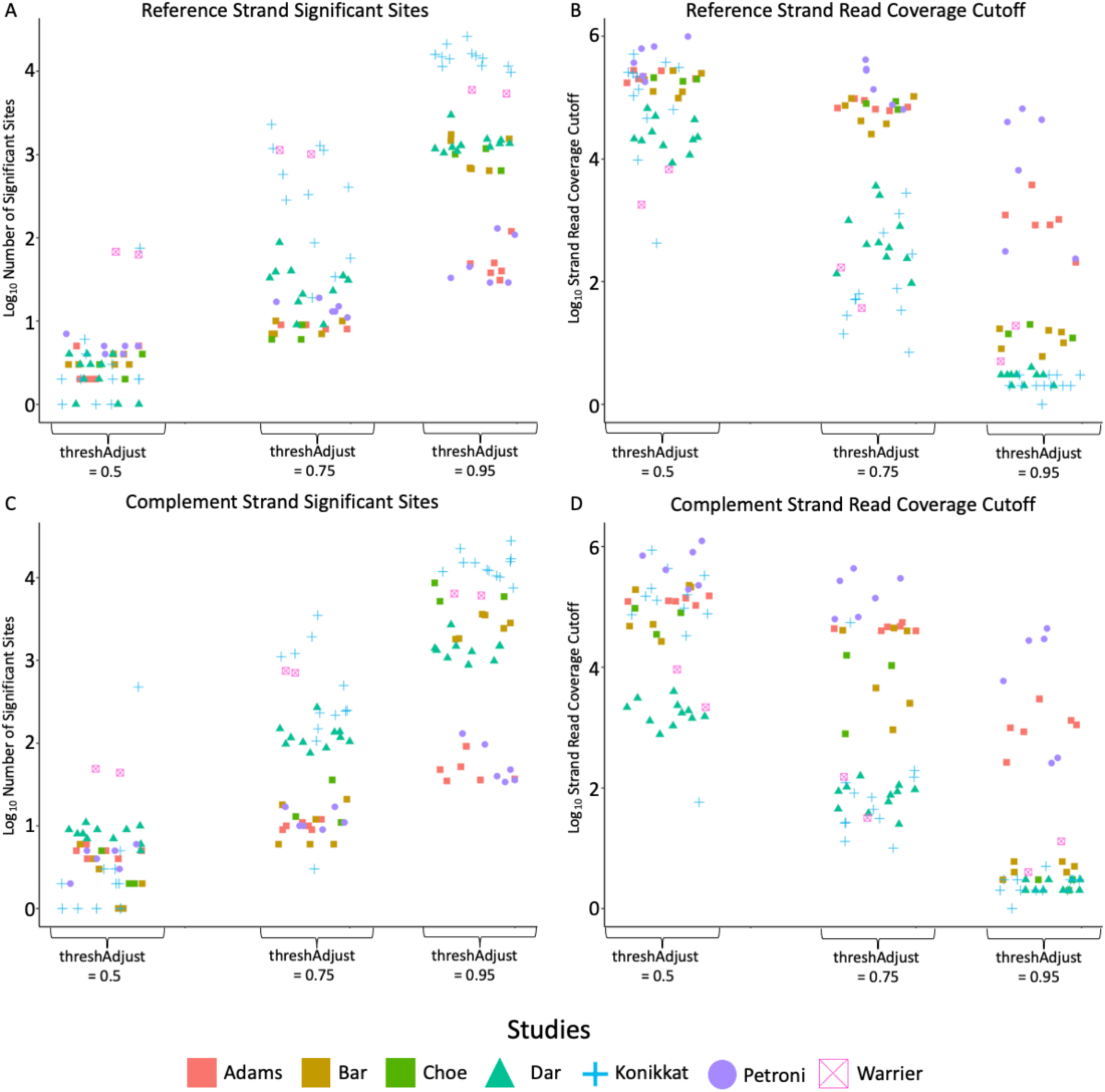
PIPETS parameterization data for the three *E. coli* studies shows different clustering patterns when compared to 3’-seq data from other organisms. A) PIPETS was run at three different threshAdjust values (highOutlierTrim = 0.01) for all three *E. coli* studies as well as 4 additional studies from four different organisms. The Dar data (mint triangle) is *B. subtilis*, the Konikkat data (blue plus) is *P. aeruginosa*, the Petroni data (purple circle) is *B. burgdorferi* and the Warrier data (pink crossed square) is *S. pneumoniae*. The log_10_ number of 3’-sites is very different at all values of threshAdjust. The Konikkat and Warrier data are consistently high outliers, but at threshAdjust = 0.95, the Bar and Choe data cluster with the Dar data and are closer to these outliers. The Adams data clusters more closely with the Petroni data. B) The reference strand read coverage cut-offs mirror the trend for sites, however the drop in cut-off value is very severe starting at threshAdjust = 0.75, but at threshAdjust = 0.95, all studies except the Adams and Petroni have similarly low values. C) The complement strand number of sites follows the same trend as the reference strand, but the difference between the Adams and Petroni samples and the rest of the studies is even greater. D) The complement strand read coverage cut-offs are even lower than the reference strand, and the distancing between the Dar, Konikkat, and Warrier studies is even greater at threshAdjust = 0.75. However, at threshAdjust = 0.95, we see that the Bar and Choe studies have lower read coverage cut-off values than the Warrier data.

## Discussion

Across the field, there are large discrepancies in analysis methodologies and results interpretation approaches which hamper the ability to derive a common consensus on the biological relevance of 3’-seq data. By using a consistent analysis and interpretation approach, we have shown that, even for 3’-seq data sets which were generated using very similar experimental conditions, extraction protocols, and sequencing techniques performed on the same strain, there are still large differences in the raw 3’-seq data as well as the potentially biologically significant results. These differences are not proportionate to changes in read depth between the files and there are many instances where changes in read coverage distribution amounts to dramatically different 3’-seq profiles between different studies (Figures 5 & 6). The inconsistencies in the 3’-seq data and results for different studies using the same organism severely limit our ability to confidently differentiate between true biological relevant changes and simply inconsistent 3’-seq data generation.

One of the most surprising findings of this work was presence of two distinct, *E. coli* 3’-seq read coverage profiles that were defined by differences in strand specific read coverage distribution. While it is accepted that some variation will always occur during experimental or sequencing protocols as a result of human or technical error, we were surprised to see such differences in read coverage patterning and distribution in these data. The Adams data had consistent 3’-seq patterning for both the reference and complement strand, unlike the large complement strand skew seen in the Bar and Choe data. Although the distribution of protein coding genes is roughly even between the reference and complement strands of *E. coli*^29^, other biological factors such as the distribution of highly expressed RNAs such as tRNA’s, 63% of which are found on the reference strand^32^, could influence the distribution of 3’-seq read coverage. While it is difficult to identify and account for all sources of variation between these studies, we do not expect that changes in protocols, even those that are not well described in available information from studies, are solely capable of creating such discrepancies. We cannot speculate the complete set of causes that resulted in the differences identified in the raw 3’-seq data of these three studies. However, we show several instances of dramatically different 3’-seq read coverage distribution and characteristics for biologically relevant sites (Figures 5 & 6) which could greatly influence any comparison done between these studies. Because each study used the same strain and very similar experimental conditions, we would expect to see much more overlap in the results and more consistent patterns of 3’-seq read coverage.

The parameter and results values we used to identify and describe differences in 3’-seq data sets are not perfect markers for fully capturing the attributes which define these *E. coli* 3’-seq data sets. The threshAdjust values for each data set that we chose for this work likely result in PIPETS dismissing biologically relevant 3’-seq signal that could influence our perception of similarity across the studies. However, we have shown in previous work^18^ and the work done here that PIPETS is consistent and robust in its ability to identify the most prominent 3’-seq signal for any data set. Furthermore, since PIPETS uses read coverage cut-offs that are generated for each data set uniquely based on the data therein, it can provide insight into the distribution of 3’-seq read coverage across a replicate. Many of the *E. coli* data sets used here were dominated by a small number of 3’ sites comprised of very high numbers of 3’-seq reads. We show that even with reasonable changes to the parameters of PIPETS, these high read coverage 3’-sites are almost always identified as significant (Figure 3).

We would expect that these high read coverage 3’-sites, which are the most likely to be derived from biologically relevant processes, would be found consistently across most sequencing results taken from the same organism. However, we did not see enough overlap between all three studies for their reported results (Figure 1) as well as the PIPETS results (Figure 3) that would suggest that the same signal from consistent biological processes is being identified. While the lack of overlap for the PIPETS results points to differences in the 3-seq data, the lack of overlap between the reported results of the three studies points to the current field standard of using study specific analysis tools to produce results. This makes it very difficult to confidently state if these different reported results differences are biological or technical differences in the data or simply the result of diverging analysis techniques and data interpretations.

We acknowledge that analyzing additional 3’-seq data sets might help to identify more consistencies compared to the pool of data used here. However, despite the growing amount of 3’-seq for both of these organisms, there are no ground-truth 3’-seq data sets for *E. coli* or any organism in the field; without which we are unable to confidently identify if additional 3’-seq data is guaranteed to provide insight in a biologically accurate representation of the population of 3’-ends of transcripts. Furthermore, we identified consistent heterogeneity in the 3’-seq data from data sets that were replicates from the same condition and study. All three *E. coli* data sets had replicates with read coverage cut-offs or significant 3’-site counts that differed from other replicates by up to an order of magnitude (Supplemental Table 5). The complement strand bias was only present in the Bar and Choe data, and the magnitude of this bias was not always consistent between replicates or studies. These replicate level inconsistencies detract from the certainty that additional 3’-seq data will always improve analytical success.

With the inconsistencies present in the raw 3’-seq data for these studies, we sought out computational methods used in the field to provide alternative context to the comparative metrics used in this work. To attempt to associate the PIPETS results with biologically relevant processes which are associated with 3’-ends, we compared the PIPETS significant 3’-sites with the predicted terminators from three different tools: RNIE^22^, RhoTermPredict^21^, and TransTermHP^23^. We found little overlap between the predicted terminators RNIE and RhoTermPredict and the PIPETS results and some overlap with TransTermHP on the complement strand (Supplemental Figure 3). This lack of correspondence further confounds our ability to confidently identify consistent, organism specific 3’-seq characteristics, making comparative studies even more challenging.

This work was performed on studies from one of the most prominent, best annotated model organisms in the field of bacterial transcription. When we look to compare *E. coli* 3’-seq data with the 3’-seq data from different model organisms, it is even more difficult to ensure that technical differences in approach are not influencing potentially biologically relevant findings. The complement vs. reference strand difference that was identified across data sets from evolutionary distant organisms is surprising as it is unlikely that different organisms with different strand specific distributions of genes, sequenced under different experimental conditions would result in such similar 3’-seq read coverage patterning. While differences in gene distribution between leading and lagging strands of DNA replication have been described^33–35^, replication is bi-directional relative to the reference strand^36–40^, and thus such differences would be expected to average out over the genome. It possible that, instead, consistent technical practices are skewing protocols to preferentially sequence and identify populations of 3’-seq read coverage that are not indicative of a given organisms true 3’-seq profile. However, given the differences in growth conditions, necessary experimental and sequencing protocols for different organisms, the biological heterogeneity between species, and any number of additional factors, it is impossible to identify any singular cause that can explain the differences and surprising similarities we identified in this work.

Finally, there are multiple different techniques currently being used in the field to generate signal from 3’-ends of transcripts. The data analyzed in this study is generated using 3’-end sequencing, which is the most direct method to produce signal for 3’-ends. Other studies use bulk-RNAseq approaches and then analyze the data with computational tools to extrapolate 3’-end signal, but often choose to directly refer to this data as “Term-seq” data. These two approaches result in the inability to directly compare raw data for many studies across the field. Further, because there are several approaches to converting RNA-seq data to 3’-seq signal, there is even less consistency in the analytical choices made for producing 3’-seq data. However, ultimately such approaches may be less susceptible to the signal/noise issue of direct 3’-end sequencing. As we have shown in this work, even 3’-seq data generated using very similar methods can have wide reaching discrepancies. We encourage the field to transition to an ongoing practice of analyzing and comparing multiple data sets using common analysis tools. Only through a consistent analytical framework can such dissimilarities be identified in data and acknowledged during interpretation of the results.

## Acknowledgements

This work was supported by grants R21AI148895 and R01GM134259 to M.M.M.

## Supplemental Tables

<u>Supplemental Table 1:</u> File identification, study association, and accession codes for each of the replicates and data sets used in this study.

<u>Supplemental Table 2:</u> Table with PIPETS parameter values for threshAdjust and highOutlierTrim along with the reference and complement strand significant 3’-site count and read coverage cut-offs associated with those values. These values and results are used as study-specific results in this work.

<u>Supplemental Table 3:</u> List of PIPETs called peaks across all three datasets used for comparisons in Figure 4.

<u>Supplemental Table 4:</u> List if 3’-termini reported by the studies assessed here^1–3^ used to create Figure 1.

<u>Supplemental Table 5:</u> PIPETS 3’-sites counts and read coverage cut-offs for each of the replicates from the three *E. coli* studies at three different values of the parameter threshAdjust (0.5, 0.75, 0.95).

<u>Supplemental Table 6:</u> The number of genomic coordinates in each replicate that contain read quality passing (minimum score 30) read coverage values that fall into 5 categories: 0 reads, 1-49 reads, 50-999 reads, 1000-9999 reads, and >= 10000 reads.

<u>Supplemental Table 7:</u> Table of file read depths for each replicate used in this study. Columns contain raw read depth, quality score passing (minimum read quality score of 30) read depth, and strand specific quality score passing read depth.

<u>Supplemental Table 8:</u> PIPETS 3’-sites counts and read coverage cut-offs for each of the replicates from all organisms used in this work in Figure 7 at three different values of the parameter threshAdjust (0.5, 0.75, 0.95).

