## Supplemental Figures for "Analysis of 3’-seq data from multiple *E. coli* studies identifies diverging results sets and raw data characteristics despite similar collection conditions"

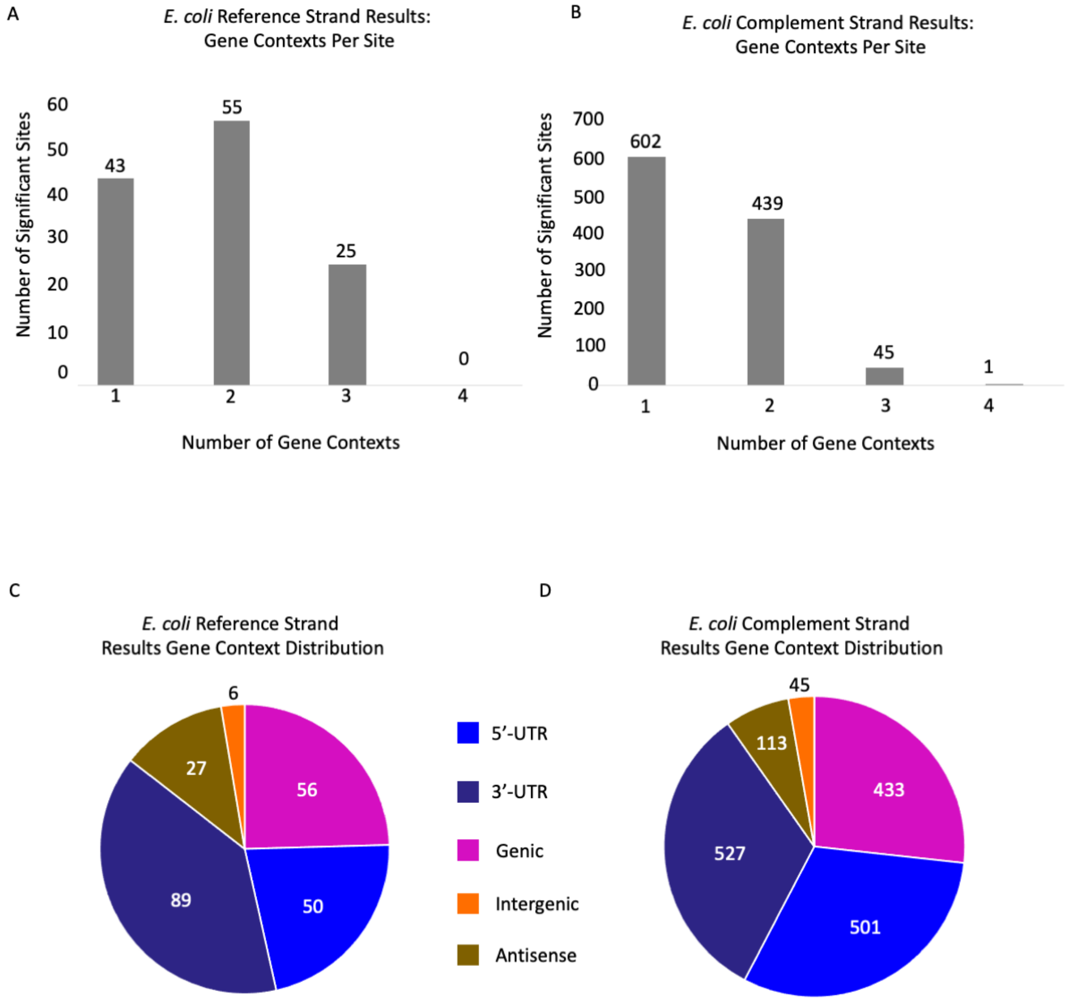


**Supplemental Figure 1: PIPETS significant 3’-sites tended to have one or two genomic contexts, which were predominantly 5’-UTR, 3’-UTR, or genic for both strands.** PIPETS results were assigned gene contexts according to their proximity to annotated genes. Gene contexts were 5’-UTR, 3’-UTR, Genic, Intergenic, and Antisense and all PIPETS results were assigned at least one of these gene contexts (Materials and Methods). A) Gene contexts per site values for all reference strand PIPETS 3’-sites (3’-sites were not filtered based on presence across studies, however duplicates sites were only counted once). Most of the reference strand *E. coli* results had one or two gene contexts per site. B) Gene contexts per site values for all complement strand PIPETS 3’-sites (same filtering process as above). Most of the complement strand *E. coli* results also had one or two gene contexts per site, but there were considerably more sites on the complement strand. C) Distribution of gene contexts for all reference strand results. If a 3’-site had multiple gene contexts, then each was accounted for individually. The largest populations of 3’-sites had 3’-UTR, 5’-UTR, and Genic contexts. D) Distribution of gene contexts for all complement strand results with the same counting method as the above. The distribution of gene contexts was comparably similar to the reference strand results, with 5’-UTR, 3’-UTR, and Genic being the largest populations.

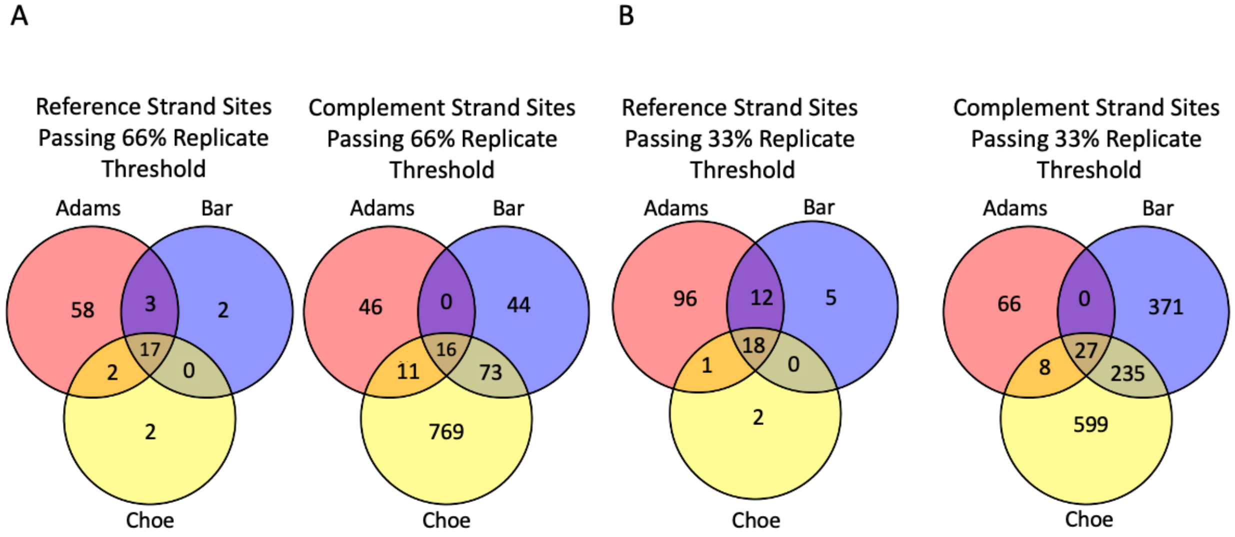


**Supplemental Figure 2: Number of PIPETS 3'-sites present after the 66% replicate threshold is lower than the number after the 33% replicate threshold, but the distribution pattern and relative overlap remains unchanged.** A) The number of 3'-sites present after the 66% replicate threshold shows very little overlap between all studies for either strand, and the Adams data is the only study with notable reference strand sites. The Bar and Choe data have many more complement strand 3'-sites with some overlap between each other, but there is almost no overlap relative to the higher number of complement 3'-sites. B) The number of 3'-sites present after the 33% replicate threshold also has very little overlap between all three studies on either strand, despite the increase in the number of complement strand sites for the Bar and Choe data. The pattern of pairwise overlap and total overlap remains the same as the 66% replicate threshold.

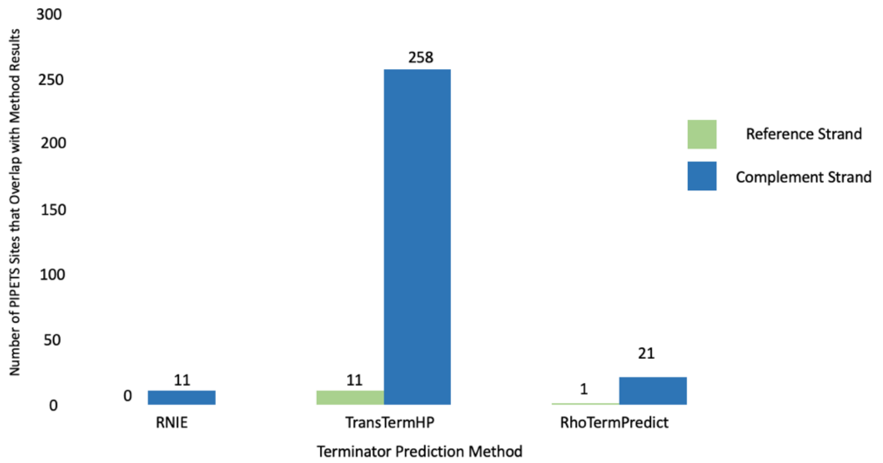


**Supplemental Figure 3: PIPETS results for both strands had limited overlap with predicted terminators from RhoTermPredict and RNIE, and some overlap with predicted terminators from TransTermHP.** Unique PIPETS results from every replicate from each *E. coli* study had little to no overlap with the predicted terminators identified by RhoTermPredict and RNIE. Only the PIPETS reference strand results had overlap with the predicted terminators from TransTermHP, but this overlap only accounts for ~25% of the total reference strand PIPETS from across all 3'-seq replicates.

**Supplemental Tables:**

Supplemental Table 1: File identification, study association, and accession codes for each of the replicates and data sets used in this study.

Supplemental Table 2: Table with PIPETS parameter values for threshAdjust and highOutlierTrim along with the reference and complement strand significant 3'-site count and read coverage cut-offs associated with those values. These values and results are used as study-specific results in this work.

Supplemental Table 3: List of PIPETs called peaks across all three datasets used for comparisons in Figure 4.

Supplemental Table 4: List if 3'-termini reported by the studies assessed here[1–3] used to create Figure 1.

Supplemental Table 5: PIPETS 3'-sites counts and read coverage cut-offs for each of the replicates from the three *E. coli* studies at three different values of the parameter threshAdjust (0.5, 0.75, 0.95).

Supplemental Table 6: The number of genomic coordinates in each replicate that contain read quality passing (minimum score 30) read coverage values that fall into 5 categories: 0 reads, 1-49 reads, 50-999 reads, 1000-9999 reads, and >= 10000 reads.

Supplemental Table 7: Table of file read depths for each replicate used in this study. Columns contain raw read depth, quality score passing (minimum read quality score of 30) read depth, and strand specific quality score passing read depth.

Supplemental Table 8: PIPETS 3'-sites counts and read coverage cut-offs for each of the replicates from all organisms used in this work in Figure 7 at three different values of the parameter threshAdjust (0.5, 0.75, 0.95).
